# Perceptual Weighting of V1 Spikes Revealed by Optogenetic White Noise Stimulation

**DOI:** 10.1101/2021.01.25.428138

**Authors:** Julian R. Day-Cooney, Jackson J. Cone, John H.R. Maunsell

## Abstract

During visually guided behaviors, mere hundreds of milliseconds can elapse between a sensory input and its associated behavioral response. How spikes occurring at different times are integrated to drive perception and action remains poorly understood. We delivered random trains of optogenetic stimulation (white noise) to excite inhibitory interneurons in V1 of mice while they performed a visual detection task. We then performed a reverse correlation analysis on the optogenetic stimuli to generate a neuronal-behavioral kernel: an unbiased, temporally-precise estimate of how suppression of V1 spiking at different moments around the onset of a visual stimulus affects detection of that stimulus. Electrophysiological recordings enabled us to capture the effects of optogenetic stimuli on V1 responsivity and revealed that the earliest stimulus-evoked spikes are preferentially weighted for guiding behavior. These data demonstrate that white noise optogenetic stimulation is a powerful tool for understanding how patterns of spiking in neuronal populations are decoded in generating perception and action.

**Significance Statement:** How the brain decodes dynamic neuronal responses to generate perception and behavior remains uncertain. A critical challenge is determining the relative contribution of spikes that occur at different times on the timescale of brain computations (tens of ms). Optogenetic tools permit causal investigations into neuronal-behavioral relationships, but are generally impractical for obtaining millisecond resolution. We circumvented this by delivering random (white noise) patterns of optogenetic inhibition to the primary visual cortex of behaving mice during visual tasks. Aligning optogenetic stimuli to task outcomes (hit, miss) yielded a neuronal-behavioral kernel – a temporal weighting that describes how inhibition at different moments impacts perception of visual stimuli. Thus, this method is a powerful tool for linking neuronal spiking, perception, and behavior.

## Introduction

How neuronal sensory signals are decoded to generate perceptions and guide behavior remains a central question in neuroscience. Quantitative measures of the relationships between sensory stimuli, population spiking, and behavior have been instrumental in guiding thinking on this subject. Measurements of correlations between neuronal responses and perceptual reports (choice probability; reviewed by Gold and Shadlen, 2007; Nienborg and Cumming, 2010; Nienborg et al., 2012; Parker and Newsome, 1998) or correlations between neuronal response latencies and reaction times (DiCarlo and Maunsell, 2005; Lee et al., 2016; Seal et al., 1983) have provided insights about which neurons are likely to contribute to a behavior, but correlative approaches require detailed assessment of noise correlations within large populations to determine which signals actually contribute to a behavior (Haefner et al., 2013), and ultimately perturbations of circuit activity are required to establish causal relationships.

Perturbing the activity of neuronal populations with electrical microstimulation or pharmacological agents can provide direct evidence of causal relationships between neuronal spiking and behavior. Optogenetic techniques have advanced such experiments by enabling spiking perturbations in genetically-defined cell types with temporal precision in the range of milliseconds (Bernstein and Boyden, 2011). However, most studies that use optogenetics to modulate spiking in behaving animals have delivered stimulation lasting hundreds of milliseconds or longer (e.g., Cone et al., 2020; Glickfeld et al., 2013; Goard et al., 2016; Guo et al., 2014; Lee et al., 2012). While extended perturbations provide valuable insights, it is important to understand how the behavioral impact of spiking of different neurons varies over intervals corresponding to the natural time scale for initiating a response to a stimulus, which for many behaviors involves only 100-200 ms (Histed et al., 2012; Sachidhanandam et al., 2013). Knowing how neurons’ spikes are weighted as a function of time during the generation of a response can reveal their relative significance to that behavior and their functional relationships to other contributing neurons. For example, Resulaj and colleagues (2018) varied in 40 ms steps the onset of a sustained optogenetic inhibition of mouse V1 neurons after the appearance of a visual stimulus and found that the first 80-100 ms of stimulus-evoked V1 activity is much more important for visual behaviors than subsequent V1 spiking. Modeling studies have suggested that the contributions of individual sensory neurons to perceptual decisions might be limited to substantially shorter intervals (Kirchner and Thorpe, 2006; Panzeri et al., 2001; Shriki et al., 2012; VanRullen and Thorpe, 2002). Optogenetic perturbations can be brief enough (<10 ms) to probe neuronal contributions with high temporal resolution (Tchumatchenko et al., 2011). However, delivering a single brief perturbation at different times on different trials can be impractical for obtaining temporal spike-weighting functions. Many trials will typically be needed to get a precise estimate of the small behavioral effects of a brief perturbation at a particular time, and many time offsets are needed to fully sample the relevant interval at high resolution. The total number of trials required will generally greatly exceed what can be obtained in a single session. Mice are the subject for most optogenetic approaches, and the high inter-session and inter-subject behavioral variability makes data from different sessions difficult to integrate.

These limitations can be overcome using the method of reverse correlation (De Boer and Kuyper, 1968). Reverse correlation using white noise stimuli has long been used as an efficient way to measure spatiotemporal receptive fields of sensory neurons (i.e., the stimulus-neuronal relationship; Aertsen and Johannesma, 1983; Dicarlo et al., 1998; Schwartz et al., 2006). When applied to behavioral reports such as a perceptual decision, rather than neuronal spiking, reverse correlation can reveal which periods of sensory stimulation dominate behavioral choices (the stimulus-behavioral relationship, Nienborg and Cumming, 2009; Okazawa et al., 2018). Here, we describe experiments in which we extended this approach by using white noise optogenetic stimulation of neuronal populations to reveal the temporal weighting of their spiking in generating a behavioral response (the neuronal-behavioral relationship). The results show that measurements of neuronal-behavioral kernels can provide a highly efficient approach to obtain spike weighting functions with high temporal resolution, and show that only very brief, stimulus dependent, epochs of V1 stimulus responses contribute to behavioral detection of visual stimuli.

## Results

### Optogenetic White Noise Stimulation of V1 During Behavior

We used transgenic mouse lines that expressed Cre-recombinase selectively in Pvalb-expressing interneurons, (PV+; Hippenmeyer et al., 2005) to restrict expression of excitatory opsins to V1 PV+ interneurons (Pfeffer et al., 2013). Each of 12 mice was implanted with a head post and a cranial window unilaterally over V1 (Goldey et al., 2014). After recovering, they were trained to do a visual detection task in which they manipulated a lever with a forepaw to report the onset of a small Gabor stimulus that appeared on a display at a random time in each trial (Cone et al., 2019; Histed et al., 2012). We then mapped V1 under the cranial window retinotopically using intrinsic signals (Figure 1A) and injected the region in V1 corresponding to the training stimulus location with Cre-dependent viruses containing ChR2-tdTomato (Nagel et al., 2003). After allowing 2-4 weeks for virus expression, an optical fiber was fixed to the implant directly above the ChR2-expressing neurons to deliver consistent optogenetic stimulation of PV+ neurons across multiple experimental sessions.

**Figure 1.**
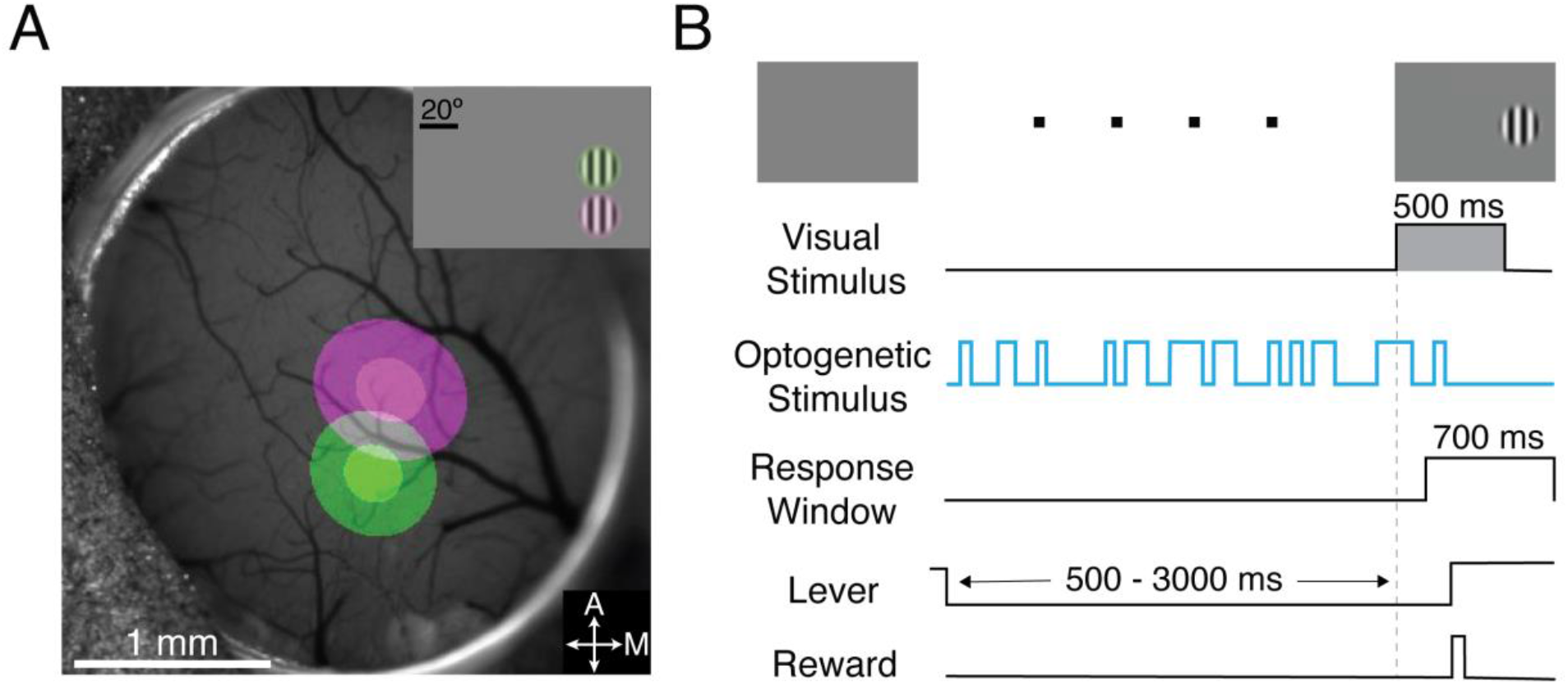
White noise optogenetic stimulation of V1 PV+ neurons during a visual detection task. **A)** Pseudo-colored intrinsic autofluorescence responses to visual stimuli presented at two visual field locations Green and magenta regions represent 2D-Gaussian fits of responses to stimuli at the different visual field locations (green: 25° azimuth, 0° elevation; magenta: 25° azimuth, −20° elevation; Gabor SD = 10°). The 1 and 2 SDs on the fits are depicted as the solid and transparent regions, respectively. A: anterior; M: medial. **B)** Trial schematic of the contrast detection task with an example white noise optogenetic stimulus profile. In some sessions the visual stimulus was a contrast step (light gray) that lasted for 500 ms, while in other sessions, the visual stimulus ramped from 0 to 100% over 500 ms.

During testing, white noise optogenetic stimulation was delivered to V1 PV+ neurons throughout one third to one half of trials (Figure 1B; see Methods). In the absence of optogenetic perturbations, mice reliably detected changes in visual contrast over many sessions (step stimulus (217 sessions): hit rate = 0.60; ramp stimulus (143 sessions): hit rate = 0.57). All remaining sections focus on trials where visual stimuli were paired with white noise optogenetic inhibition. Behavioral outcomes on trials with white noise optogenetic stimulation were used to construct a neuronal-behavior kernel (NBK; see Methods). The average effect of the optogenetic perturbation on detection performance was modest (step stimulus: median Δhit rate −0.09, IQR −0.03 to −0.17; ramp stimulus: median Δhit rate −0.07, IQR −0.03 to −0.13). For the primary analysis, optogenetic stimulus profiles preceding hits or misses were aligned with the onset of visual stimuli. Next, we normalized the high binary optogenetic power for each trace to 1 to account for the different powers used across mice with different levels of virus express (median = 0.15 mW, IQR = 0.09 - 0.25 mW). Optogenetic stimulus profiles from miss trials were inverted and then averaged together with the profiles from hit trials to generate a first-order Weiner kernel. The resulting kernel represents the linear temporal weighting that describes how optogenetically activating V1 PV+ neurons at times relative to the visual stimulus onset impacts behavioral detection of that stimulus (de Boer and Kuyper, 1968).

Figure 2A shows the result from an individual behavioral session that yielded a particularly clear NBK. A value of 0 normalized power corresponds to the presence or absence of the optogenetic stimulus having no net effect on the animal’s detection of the Gabor. The large negative peak in the neuronal-behavioral kernel ~50 ms after visual stimulus onset corresponds to reduced optogenetic stimulation at this time making the animal more likely to detect the visual stimulus. A negative peak is expected because lower optogenetic power would reduce inhibition onto principal neurons in V1. PV+ neuron stimulation before the visual stimulus or 100 ms after visual stimulus onset had little effect on behavior, even though the visual stimulus stayed on the display until the end of the trial, suggesting that the earliest period V1 spiking is most critical. This critical window of V1 activity ended ~200 ms before the animal’s responses (Figure 2A, lower histogram shows reaction times).

**Figure 2.**
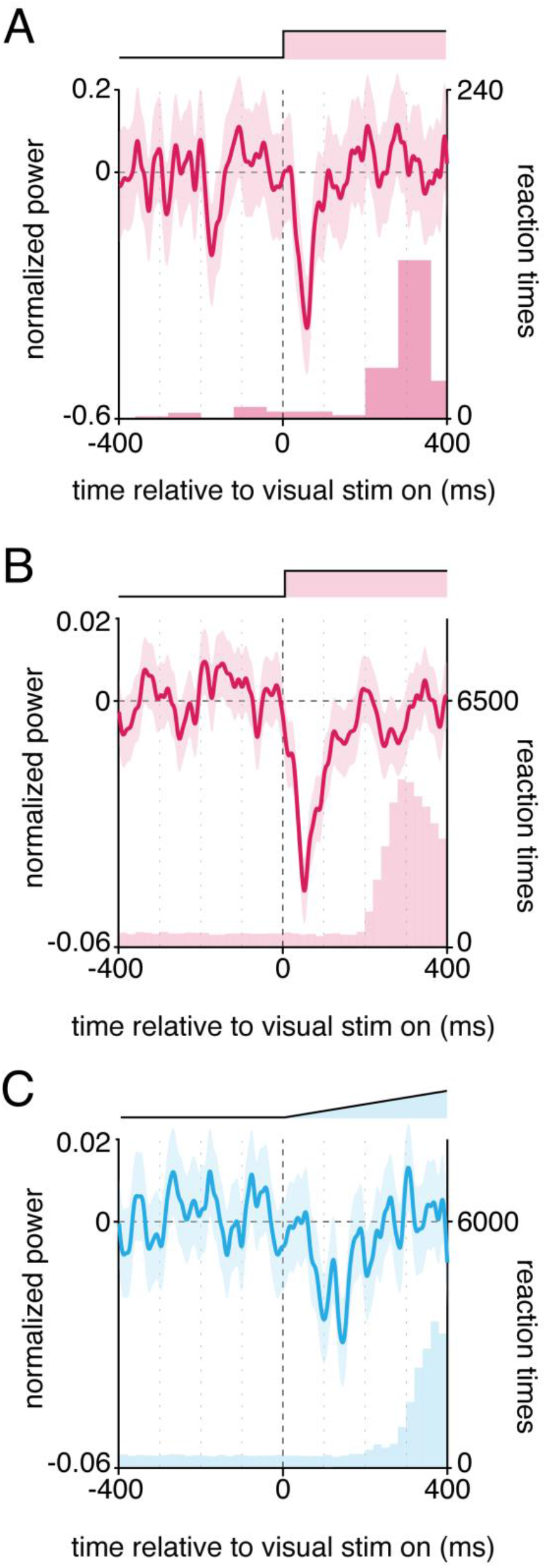
Neuronal-behavioral kernels identify the critical moments of V1 spiking underlying behavior in stimulus dependent fashion. **A)** The line represents the mean of all optogenetic stimulus profiles (hits, inverted miss profiles) normalized and aligned to the onset of visual contrast steps for a single representative behavioral session. The shaded area represents the 95% CI based on a bootstrap procedure. The profile of the visual stimulus contrast step is depicted above. Lower right histogram is the reaction time distribution. **B)** Same as in A but averaged across 33452 trials from 216 behavioral sessions from 10 mice. **C)** Same as in B, except for sessions in which the visual contrast ramped from 0-100% over 500 ms (18939 trials, 143 sessions, 6 mice). See also Figure S1-3.

Significant NBKs were not obvious in most individual sessions (which included 100-300 trials), but observations from different sessions and animals can be combined. The red trace in Figure 2B shows an average NBK based on 216 sessions from 10 mice (see Figure S1 for NBKs from individual animals). This average NBK is only slightly broader than the NBK from the single session in Figure 2A, implying that the timing of effects is relatively consistent across animals and sessions. The negative peak of the average NBK occurs 52 ms after the onset of the visual stimulus and the NBK was continuously negative from 5 to 122 ms.

In constructing the NBKs in Figure 2, optogenetic stimuli were aligned to the onset of the visual stimuli (t = 0). We used this alignment because we expected the V1 neuronal response and its behavioral contribution to be linked to the visual stimulus onset. However, other alignments are possible. In particular, the NBK can be calculated using optogenetic stimuli that have been aligned with the response time for each hit trial. A response-time-aligned NBK would be a natural choice if optogenetic perturbations were delivered to motoneurons. The response-time-aligned V1 NBK (Figure S2) was much broader than the visual-stimulus-aligned NBK in Figure 2B, suggesting that the spikes in V1 that drive the detection of a visual stimulus are those that occur time-locked to the onset of that stimulus, and that most response time variability occurs in structures that lie between the V1 neurons that contribute to the behavior and the motoneurons that generate the response (Lee et al., 2016). Other alignments, such as stretching time on individual trials to align both the stimulus onset and the response time across trials also produced a broader NBK (Figure S2). A V1 kernel constructed using optogenetic stimuli aligned to lever releases on trials with a false alarm revealed no significant peaks (Figure S2). This suggests that false alarms were not driven by fluctuations in V1 activity that caused the animals to perceive a fictive visual stimulus, but rather were driven by activity in other brain regions. This is consistent with behavioral data showing that mice cannot detect isolated activation of their V1 PV+ neurons (Cone et al., 2019, 2020). Control measurements made with optogenetic stimulation that was misaligned with the opsin-expressing neurons showed that the behavioral effects did not arise from cortical heating, direct retina stimulation by the optogenetic stimulus, or non-specific behavioral effects (Figure S3).

The relationship between neuronal activity and behavior revealed by an optogenetic kernel will be specific to for the population of neurons stimulated (e.g., sensory and motor neurons will contribute at different times), the stimulus used (e.g., perturbation of V1 neurons might have less impact on detecting brightness changes than contrast changes) and task (e.g., an animal presented with simultaneous auditory and visual stimuli should show different effects from V1 perturbations depending on whether the animal is tasked with discriminating visual or auditory stimuli). To see whether different neuronal-behavioral kernels can be readily resolved, we optogenetically stimulated V1 in some of the same mice while they detected a different visual stimulus that required longer integration. We conducted separate sessions in which the contrast of the Gabor stimulus ramped linearly from 0% to ~100% contrast over 500 ms, while keeping all other stimulus and task parameters the same.

As expected, the median response time for the ramping stimulus was longer than for contrast steps (by 63 ms: 413 ms versus 350 ms; lower histograms in Figure 2B, C). The negative peak of the NBK was delayed for ramping Gabors relative to stepped Gabors (blue trace, Figure 2C). The peak of the average NBK was noisy, but reached its lowest point 142 ms after the onset of the visual stimulus, with the 95% confidence interval (almost) continuously negative from 76 to 163 ms. These results show that neuronal-behavioral kernels derived from optogenetic stimulation can reveal small differences in the timing of contributions from a given neuronal population to different behaviors.

### Comparing Neuronal Behavioral Kernels and Neuronal Activity in V1

The early occurrence of the neuronal-behavioral peak points to behavior depending on the spikes that occur immediately following stimulus onset. However, the timing of the kernel cannot be compared directly with spiking in V1 because the effects of optogenetic stimulation on V1 spiking will be delayed by the latency of PV cell spiking in response to optogenetic stimulation and delays in the propagation of inhibitory signals to principal cells (Packer and Yuste, 2011). There will also be polysynaptic, network effects that will unfold in V1 over a longer time course (Li et al., 2019). Fortunately, all of the relevant delays can be captured and addressed by recording the responses of V1 neurons to optogenetic white noise stimulation. A spike-triggered average (STA) of a white noise stimulus provides the impulse response of a neuron (Bryant and Segundo, 1976) and the aggregated STA of V1 neurons provides the overall response of V1 spiking to an impulse of optogenetic PV stimulation and captures the relevant delays in changes in spiking.

We made acute electrophysiological recordings from single and multiunit sites in V1 using multielectrode arrays in awake mice (n=5) expressing ChR2 in PV+ neurons, collecting responses to stepped or ramped Gabor stimuli in different data sets (see Methods). Visual responses from mouse V1 are characteristically weak (Glickfeld et al, 2013; Seigle, et al, 2019), and because we recorded simultaneously from many neurons, the stimuli were not optimal for most units. When the stepped Gabor was presented, 38% of units (24/63) were significantly excited and 3% (2/63) were significantly inhibited (p < 0.05, Wilcoxon signed-rank test). The median response across all units was 0.6 spikes/s (p < 10^-6^, Wilcoxon signed-rank test). A different set of neurons was tested with ramped Gabors, for which 49% of units (34/70) were significantly excited and 14% (10/70) were significantly inhibited (p < 0.05, Wilcoxon signed-rank test). The median response to ramped Gabors across all units was 0.4 spikes/s (p < 10^-5^, Wilcoxon signed-rank test). On a fraction of trials, a white noise optogenetic stimulus was delivered before, during and after the Gabor, using a power in the range used in the behavioral studies (0.25 mW mean power). Among individual units that were significantly modulated by optogenetic stimulation, most were inhibited (stepped Gabors: 22/33 (66%) inhibited, 11/33 (33%) excited; ramped Gabors: 40/53 (75%) inhibited, 13/53 (25%) excited, all p < 0.05, see Methods), but the low power of the optogenetic stimulus caused only a modest reduction in spiking (stepped Gabors: n = 63; median −0.14 spikes/s, IQR −0.42 to +0.10; ramped Gabors: n = 70; median −0.27 spikes/s, IQR −1.3 to +0.10; both p < 0.05, both Wilcoxon signed-rank test).

The STA is a reverse correlation of the optogenetic stimuli aligned to the occurrence of individual spikes (at t = 0). We measured significant STAs in 41/70 units that were tested (see Methods for inclusion criteria). The population STAs shown in Figures 3A,B thus reveal the average delays between optogenetic stimulation delivery and changes in spiking. Three of the V1 units that were excited by optogenetic stimulation appeared to be PV+ cells expressing ChR2, as their STAs showed they spiked with short latencies (5-10 ms) following positive deflections in the optogenetic stimulus power. The aggregated STA for these units is plotted in Figure 3A. The STA has a large positive peak 2 ms before spikes occurred, showing that putative PV+ cells responded with very short latency following a step in the optogenetic stimulus power. This STA peak is artifactually widened by the pulse width that we used for the optogenetic stimulus, which broadened peaks by 25 ms (which is why the positive peak extends to times after the spike, when optogenetic stimulation could have no influence on that spike).

**Figure 3.**
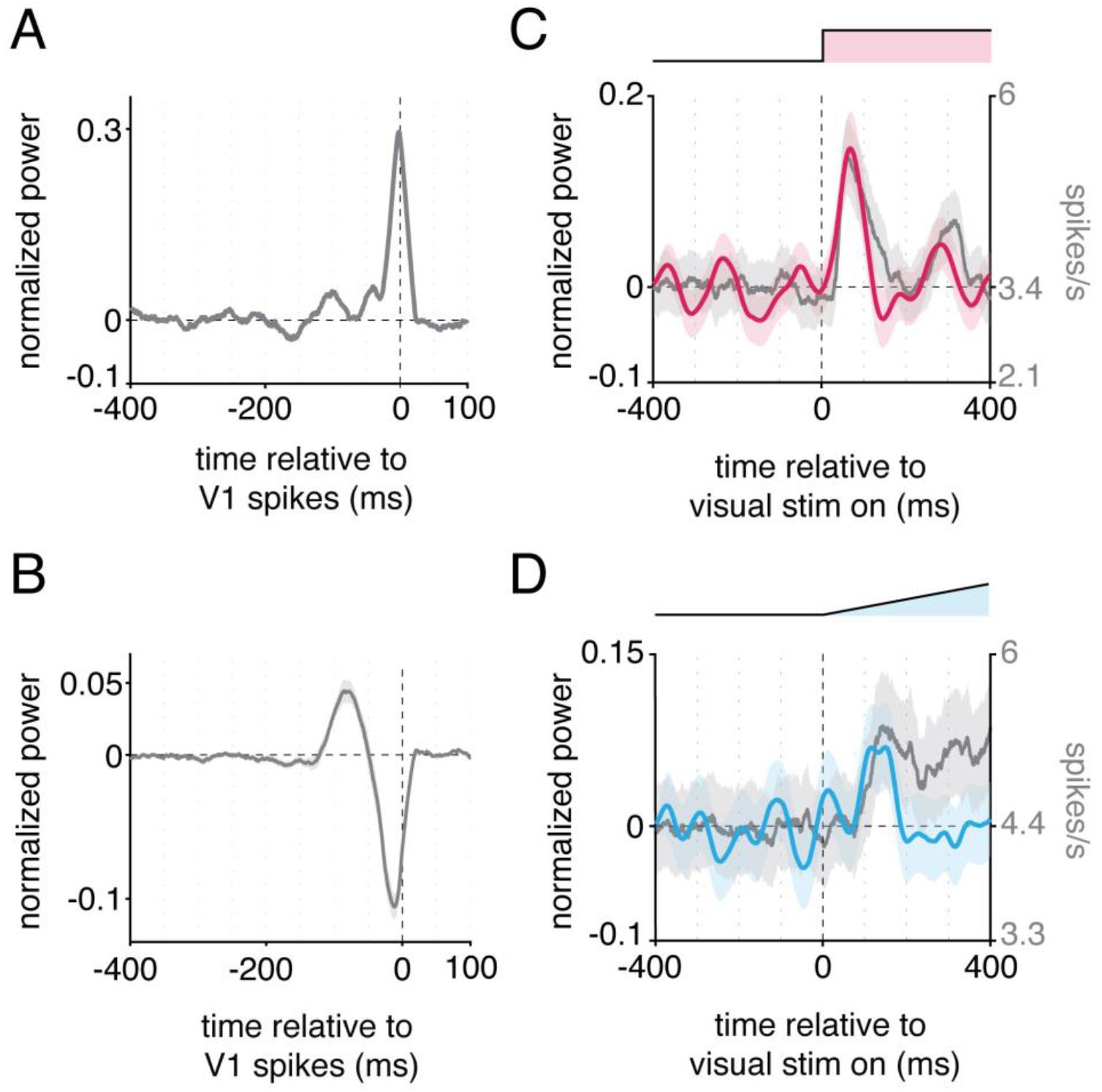
V1 spike-triggered average optogenetic stimuli and delay-corrected effects of neuronal-behavioral kernels on V1 spiking. Optogenetic stimulus profiles before and after V1 spikes were aligned and averaged for each recorded V1 unit to calculate a spike-triggered average (STA) optogenetic stimulus. Only cells that met inclusion criteria are shown (41 of 70 units, see Methods). **A)** The gray trace depicts the STA averaged from three putative PV+ units. Spikes immediately followed positive deflections in optogenetic stimuli. **B)** Same as in A, except for the remaining, putative-principal V1 cells (n=38). **C)** Colored trace (left y-axis) represents the convolution between the average neuronal-behavioral kernel measured for contrast steps (Figure 2B) and the population STA for principal neurons. The gray trace represents the population PSTH (n=63 units; right y-axis). **D)** Same as in C, except for Gabor contrast ramps (blue; PSTH = 70 units). Shaded bands are 95% CIs (convolutions) or SEMs.

The average STA for the remaining V1 units (n = 38) that were not directly excited by optogenetic stimulation (e.g., putative principal neurons) is plotted in Figure 3B and has a distinctly different form. It shows that most V1 neurons tend to spike ~12 ms after a drop in optogenetic power. An earlier, positive peak implies there is a rebound effect such that spiking is more likely if PV stimulation was stronger than average ~80 ms in the past. In particular, the timing of the negative peak of the STA indicates that the dominant effects of optogenetic PV stimulation (such as the NBK) on overall V1 spiking (the PSTH) are delayed by 12 ms.

With the assumption of linearity, we can directly compare the effect of the NBK on V1 spiking by convolving the population STA (the average impulse response function of V1) with the NBK. We focus on the STA for the putative non-PV+ cells, which include the cells that send signals to the rest of the brain. The convolution captures all the delayed changes in spike rate (both positive and negative) to optogenetic stimulation. Figure 3C shows the convolution for the stepped Gabor kernel (red) superimposed on the average response (PSTH) of those cells when no optogenetic stimulation was applied (gray). Figure 3D shows the corresponding plots for the ramped Gabor stimulus (blue and gray). As expected, responses to the stepped Gabor were earlier and stronger than responses to the ramped Gabor. Because they account for delays, the peaks of the convolutions are somewhat later than those for the kernels in Figure 2B,C (68 ms versus 52 ms for stepped, 150 versus 145 ms for ramped, although the ramped peak is noisy) and close to the time of the peaks in the PSTHs (60 and 142 ms). In both cases, the convolutions appear to return to baseline more rapidly than the PSTHs, indicating that the earlier portion of the V1 response contains the spikes that contribute most to the behavioral response. Spikes in the later portion of the responses appear to have far less weight in driving behavioral responses.

## Discussion

White noise sensory stimuli have long been used to study information transfer across multiple stages in the nervous system (Marmarelis and Naka, 1972). White noise optogenetic stimulation has recently been used to examine neuronal contributions in invertebrates as they engaged in natural behaviors, but with temporal modulations at 4 Hz or slower (Hernandez-Nunez et al., 2015; Porto et al., 2019). We took advantage of the fast and potent inhibition PV+ interneurons exert on local neurons (Packer and Yuste, 2011) to use optogenetic white noise stimulation to test which spikes in mouse V1 are most important for driving detection of a visual stimulus. We obtained neuronal-behavioral kernels showing that only spikes occurring within the first ~100 ms of a V1 response contribute strongly to detection. Our results are consistent with previous experimental work using steps of opsin stimulation (Resulaj et al., 2018), but provide a complete spike weighting function that is corrected for the delays inherent in optogenetic stimulation.

The outsized importance of the earliest stimulus-evoked spikes in primary sensory areas has long been inferred based on the presence of fully-developed, strong stimulus selectivity at the very start of neuronal responses and complementary modeling studies examining the performance of early spikes (Bair, 1999; Chase and Young, 2007; Tovée, 1994; VanRullen and Thorpe, 2002). Dependence on a relatively small proportion of early spikes in a sensory response is consistent with visual capabilities such as fine discrimination of faces that are masked after ~50 ms of viewing (Lehky, 2000) and responses in demanding visual categorization tasks being completed in ~120 ms (Kirchner and Thorpe, 2006). Other studies have similarly pointed toward the earliest spikes being preferentially used in a variety of tasks and areas (Chen et al., 2008; Müller et al., 2001; Oram and Perrett, 1992). Shriki and colleagues (2012) found that visual stimulus orientation could be decoded using only the timing of the first spikes from small populations of neurons. Given the wide range of response latencies seen for neurons within individual cortical visual areas (Schmolesky et al., 1998), neurons with the longest response latencies (typically near the surface of cortex: Best et al., 1986; Maunsell and Gibson, 1992; Raiguel et al., 1999) might contribute relatively little to fast behavioral detection.

It is important to recognize that the kernels we measured represent upper bounds on the intervals over which V1 spikes drive behavior. The 25 ms step size of our optogenetic stimulus artificially broadens the kernels and STAs. The peaks in our plots could correspond to actual functions that are as much as 50 ms narrower at their base. Future experiments could address this potential broadening by using opsins with faster dynamics (e.g., ChETA, Chronos; Gunaydin et al., 2010; Klapoetke et al., 2014) although no existing opsins approach 1 ms resolution. Alternatively, substantially larger data sets might support analytical methods that compensate for the broadening (e.g., deconvolution).

White noise optogenetic stimulation offers many advantages for exploring how specific brain structures and cell types contribute to behavior. Because it combines trials across many sessions and animals, relatively weak stimulation can be used, allowing the brain to remain close to its natural operating state. Because the stimulus is present throughout the trial, it efficiently samples the entire perceptual/behavioral cycle in an unbiased way. White noise stimulation can also simultaneously capture the full dynamics of the neuronal contributions, from DC to the limits imposed by the Nyquist frequency of the white noise.

Because each NBK is specific to a particular task (i.e., stimulus-response combination), the presence, timing and magnitude of a kernel can provide insights into whether and when particular neurons contribute during the execution of particular behaviors. The relative timing of kernels in different brain structures during a given task can provide direct evidence regarding functional relationships between spiking in different circuit elements. Even if kernels span tens of milliseconds and their breadth is limited by opsin and circuit dynamics, the timing of the peaks or centers of mass of kernels in different structures might be distinguished with millisecond precision, which would easily serve for detecting the offsets likely to be associated with the ~5 ms increase in neuronal response latencies seen between successive levels of cortical processing (Mitzdorf and Singer, 1979). These and other applications suggest that white noise neuronal-behavioral kernels can provide a powerful tool for exploring circuit-level contributions to behavior.

## Methods

### Animal Preparation

All animal procedures followed NIH guidelines and were approved by the Institutional Animal Care and Use Committee of the University of Chicago. We used mice (12 mice, 7 female) that were heterozygous for Cre recombinase in Parvalbumin (PV) expressing cells, which allows targeting specificity of ~93% (Pfeffer et al., 2013). Animals were outbred by crossing homozygous Cre-expressing PV-Cre mice (JAX Stock #017320; Hippenmeyer et al., 2005) with wild-type BALB/c mice (JAX stock #000651). Animals were singly housed on a reverse light/dark cycle with *ad libitum* access to food. Mice were water scheduled throughout behavioral experiments, except for periods around surgeries.

Mice (3–5 months old) were implanted with a headpost and cranial window over V1 to give stable access for photostimulation during behavior (Goldey et al., 2014; Histed and Maunsell, 2014). For surgery, animals were anesthetized with isoflurane (induction, 3%; maintenance 1-1.5%) and given ketamine (40 mg/kg, IP) and xylazine (2 mg/kg, IP). Body temperature was maintained with a regulated heating pad. A titanium headpost was secured to the skull using acrylic (C&B Metabond, Parkell) using aseptic technique. A craniotomy was made over V1 in the left cerebral hemisphere (3.0 mm lateral and 0.5 mm anterior to lambda) and covered with a glass window (3.0 mm diameter, 0.8 mm thick; Tower Optical). Following surgery, mice were given analgesics (buprenorphine, 0.1 mg/kg and meloxicam, 2 mg/kg, IP).

After recovery from surgery, we located V1 by measuring changes in the intrinsic autofluorescence signal using visual stimuli and epifluorescence imaging (Andermann et al., 2011). Autofluorescence produced by blue excitation (470 ± 40 nm, Chroma) was collected using a green long-pass filter (500 nm cutoff) and a 1.0X air objective (Carl Zeiss; StereoDiscovery V8 microscope; ~0.11 NA). Fluorescence was captured with a CCD camera (AxioCam MRm, Carl Zeiss; 460 x 344 pixels; 4 x 3 mm FOV). The visual stimuli were full contrast drifting Gabors (10° SD; 30°/s; 0.1 cycles/deg) presented for 10 s followed by 6 s of mean luminance at multiple visual field locations. The response to visual stimuli was computed as the fractional change in fluorescence during the first 8 s of the stimulus presentation compared with the average of the last 4 s of the preceding blank.

Virus injections were targeted to the monocular region of V1 based on each animal’s retinotopic map (25° azimuth; ±15° elevation). Mice were anesthetized (isoflurane, 1%–1.5%), their head post was secured, and the cranial window was removed using aseptic technique. We used a volume injection system (World Precision Instruments) to inject 200–400 nl of AAV9-Flex-ChR2-tdTomato (~10^11^ viral particles; Addgene) 250-400 μm below the cortical surface (50 nl/min). Following the injection, a new cranial window was sealed in place. Two to three weeks after injection, we localized the area of ChR2 expression using tdTomato fluorescence, and attached an optical fiber (400 μm diameter; 0.48 NA; Doric Lenses) within 500 μm of the cranial window (~1.3 mm above the cortex).

### Behavioral Task

Mice were trained to use a lever to respond to changes in a visual display for a water reward (Histed et al., 2012). During behavioral sessions, mice sat comfortably in a sled that enabled head fixation. Mice were positioned in front of a calibrated visual display that displayed a uniform gray field. Mice self-initiated trials by depressing the lever and a neutral tone indicated the start of each trial. The prestimulus period was randomly drawn from a uniform distribution (500-3000 ms), after which an achromatic, odd-symmetric Gabor stimulus (5° SD, 0.1 cycles/deg) appeared. Mice had to release the lever within 100-700 ms after stimulus onset (hit) to receive a liquid reward (1.5-4 μL). Trials in which mice failed to release the lever within the response window (miss) resulted in a brief time out (1500-2500 ms). Trials in which mice released the lever before the onset of visual stimuli were counted as false alarms and were excluded from all analyses (except Figure S2). Task parameters were slowly adjusted over several weeks until mice responded reliably to Gabor stimuli spanning a range of contrasts in the location of the V1 retinotopic map where optogenetic stimulation would be delivered during testing sessions (>60% hit rate for a range of stimulus contrasts, >300 trials a day; median training time: 61 days; IQR: 48-62 days). For sessions with contrast ramps, the same Gabor (5° S.D., 0.1 cycles/deg) appeared, but the contrast increased linearly from 0% to ~100% over 500 ms.

### White Noise Optogenetic Stimulation and Analysis

Optogenetic stimulation began >4 weeks after ChR2 injection. Light was delivered through the optic fiber using a calibrated LED source (455 nm; ThorLabs). To prevent mice from seeing optogenetic stimuli, we enclosed the fiber implant in blackout fabric (Thor Labs) attached to the head post using a custom mount. The optogenetic stimulus delivered on each trial was synchronized with the visual stimulus using a photodiode mounted on the display. To avoid an abrupt onset to the optogenetic stimulus, its power ramped up to the average mean power over the first 250 ms of stimulation.

In preliminary sessions, the optogenetic stimulus for each animal was adjusted by presenting a constant stimulus power throughout trials. These sessions were used to set a power that produced a small decrease in the proportion of correct trials (~5-10% reduction in the proportion correct), which was then used as the mean power in subsequent testing sessions. The selected power varied between mice, likely owing to differences in the strength and spatial distribution of virus expression, optic fiber alignment, and behavioral capacity (median: 0.15 mW, IQR 0.09 - 0.25 mW). Preliminary sessions used to determine stimulation powers were not included in analyses.

For trials with optogenetic stimulation, binary white noise optogenetic stimuli were generated by randomly assigning zero or maximum (2x mean) power to a series of 25 ms bins with equal probability. Substantially briefer bins could not accommodate the kinetics of the ChR2 (~10 ms decay time; Mattis et al., 2012) as well as the kinetics of the PV-to-principal neuron synapse (~15 ms rise and decay time; Packer and Yuste, 2011). The phase of the 25 ms bin sequence was randomly offset to the nearest millisecond on each trial. The resulting optogenetic stimulus produced equal power across all frequencies represented and is, therefore, a quasi-white noise stimulus (Marmarelis and Marmarelis, 1978). Optogenetic stimulation was delivered on a random subset of trials (33-50%). Optogenetic stimulation was only delivered in sessions in which the initial performance without stimulation had false alarm rates < 40% and hit rates > 60% (568 of 699 sessions).

A first-order Weiner kernel was calculated from the optogenetic stimuli for trials that ended with a hit or a miss. Data from different animals were combined by normalizing the high level of the binary optogenetic power to 1. For the primary analysis, the optogenetic stimulus profiles from hit and miss trials were aligned to the onset of the visual stimulus, profiles from miss trials were inverted, and then averaged for all trials (14,612 hit trials, 15,811 miss trials for the stepped contrast stimulus; 8,984 hit trials, 10,076 miss trials for the ramped contrast stimulus). Kernels were then low-pass filtered with a corner frequency of 90 Hz. The resulting neuronal-behavioral kernel represents the linear temporal weighting of optogenetic activation of V1 PV interneurons for perturbing visual contrast detection.

In a separate analysis, we computed an NBK after aligning the optogenetic stimuli to the onset of behavioral responses (Figure S2), which would reveal neuronal contributions that were temporally aligned with the response. We also computed an NBK after stretching the time between stimulus onset and reaction time in each trial to a fixed interval, which would reveal neuronal contributions that occurred midway between stimulus onset and behavioral response. Finally, we computed an NBK using optogenetic stimuli aligned to false alarms, to determine whether those errors could be associated with optogenetically driven fluctuations in spike in V1.

In control sessions, the visual stimuli were offset from the visual field representation at the V1 site of optogenetic inhibition (Figure S3), using 3 mice previously tested with ramped Gabors. After kernels were obtained on 5 consecutive training days with visual and optogenetic stimuli aligned, mice were tested with the same stimulus presented in the opposite hemifield for 5-10 sessions in each subject (Figure S3A). For two of the three mice, the visual stimulus was returned to the original aligned configuration and additional data were collected (Figure S3B).

### Electrophysiological recordings

We recorded extracellularly from V1 in awake, passively viewing, head fixed mice using multisite silicon probes (Neuronexus; 32-site model 4 × 8-100–200-177) in 10 recording sessions in five mice (1 female). All mice were experimentally naive. Prior to recording sessions, mice were surgically prepared with headposts and optical windows, mapped for retinotopy, and injected with ChR2 as described above.

Electrophysiology sessions occurred >4 weeks after injection. electrodes were electroplated with a gold solution mixed with carbon nanotubes (Ferguson et al., 2009; Keefer et al., 2008) to impedances between 200-500 kΩ. Mice were anesthetized with isoflurane (1.2–2% in 100% O2) and head fixed. A gamma corrected visual display was positioned in the visual hemifield opposite of the recording site (~10 cm viewing distance). The eyes were kept moist with 0.9% saline throughout the session. We visualized ChR2-expressing areas of V1 by imaging tdTomato fluorescence with a fluorescence microscope and camera (Zeiss). The cranial window was then removed and the electrodes lowered through a slit in the dura. We then positioned an optic fiber above the cortex at a distance comparable to that used during behavioral experiments (1.0-1.5 mm). The craniotomy was then covered with 3% agar (MilliporeSigma) dissolved in aCSF (TOCRIS). The agar was prevented from drying out by covering it with silicone oil. After an hour recovery, anesthetic was removed and we waited at least an additional hour before recording.

The electrode was advanced in V1 and was allowed to settle for 30 minutes before collecting data. Delivery of stimuli and data acquisition were computer controlled. Concurrent visual and optogenetic stimuli were similar to those used during behavioral experiments (stepped contrast stimuli: a Gabor patch with SD 13°; ramped contrast stimuli: full screen grating 500 ms duration). We recorded >100 repetitions of each stimulus condition (visual stimulus, visual stimulus + white noise optogenetic stimuli).

Electrode signals were amplified, bandpass filtered (750 Hz to 7.5 kHz) sampled around amplitude threshold crossings (Blackrock, Inc.) and spikes were sorted offline (OfflineSorter, Plexon, Inc.). Analyses were done using MATLAB (MathWorks) and custom code. We recorded both single and multi-units but did not differentiate between them because our primary interest was how optogenetic manipulations affected the V1 population.

### Statistical Analyses

The behavioral analysis included only mice produced at least 8 days of behavioral data while being optogenetically stimulated. Because activation of V1 PV+ neurons reliably impairs visual perception (Cone et al., 2019, 2020; Glickfeld et al., 2013; Jin and Glickfeld, 2019; Lee et al., 2012), only behavioral sessions with a lower proportion of correct responses on optogenetic stimulation trials compared to unstimulated trials were used. The confidence intervals were generated using a bootstrap procedure with 10,000 draws with replacement from the measure being tested.

For electrophysiology, units were considered visually responsive to contrast steps (without optogenetic stimulation) if they exhibited a significant change in their average firing rate (p < 0.05; Wilcoxon signed-rank test) during the 50-150 ms after stimulus onset relative to the average firing rate during the 50-150 ms before stimulus onset. For the ramped contrast stimulus, the analysis window extended to the end of visual stimulus presentation (50-500 ms), relative to 50-500 ms before visual stimulus onset. Units were defined as optogenetically responsive if there was a significant difference (p < 0.05; Wilcoxon signed-rank test) in the average firing rate 50-500 ms after visual stimulus onset between trials with and without optogenetic stimulation.

We calculated the spike triggered average (STA) optogenetic stimulus based on standard approaches (Schwartz et al., 2006). The power used for binary optogenetic stimuli was first normalized from −1 to 1. For each unit, optogenetic stimuli in windows surrounding 350 before to 100 ms after individual spikes occurred were extracted and averaged. Only units for which the minimum or maximum of the spike triggered average exceeded the 95% confidence intervals by a factor of 4 (~8 SEM) were used in further analyses (e.g., population STA, cross-correlation between neuronal-behavioral kernel and population STA). Eliminating the inclusion criteria did not qualitatively impact our observations. For convolving the kernels and the population STA, the population STA and kernel were both centered on t=0.

## Acknowledgements

We thank Dr. Supriya Ghosh, Dr. Nicolas Masse, and Matthew Rosen for helpful comments on drafts of this manuscript. This work was supported by NIH U19 NS107464 and European Research Council Project 647725 (JHRM) and a postdoctoral fellowship from the Arnold and Mabel Beckman Foundation (JJC).

## Author Contributions

JRD-C, JJC, and JHRM designed experiments, JRD-C. and JJC collected data, JRD-C., JJC, and JHRM analyzed data, JRD-C, JJC, and JHRM wrote the manuscript.

## Supplemental Figures

**Figure S1.**
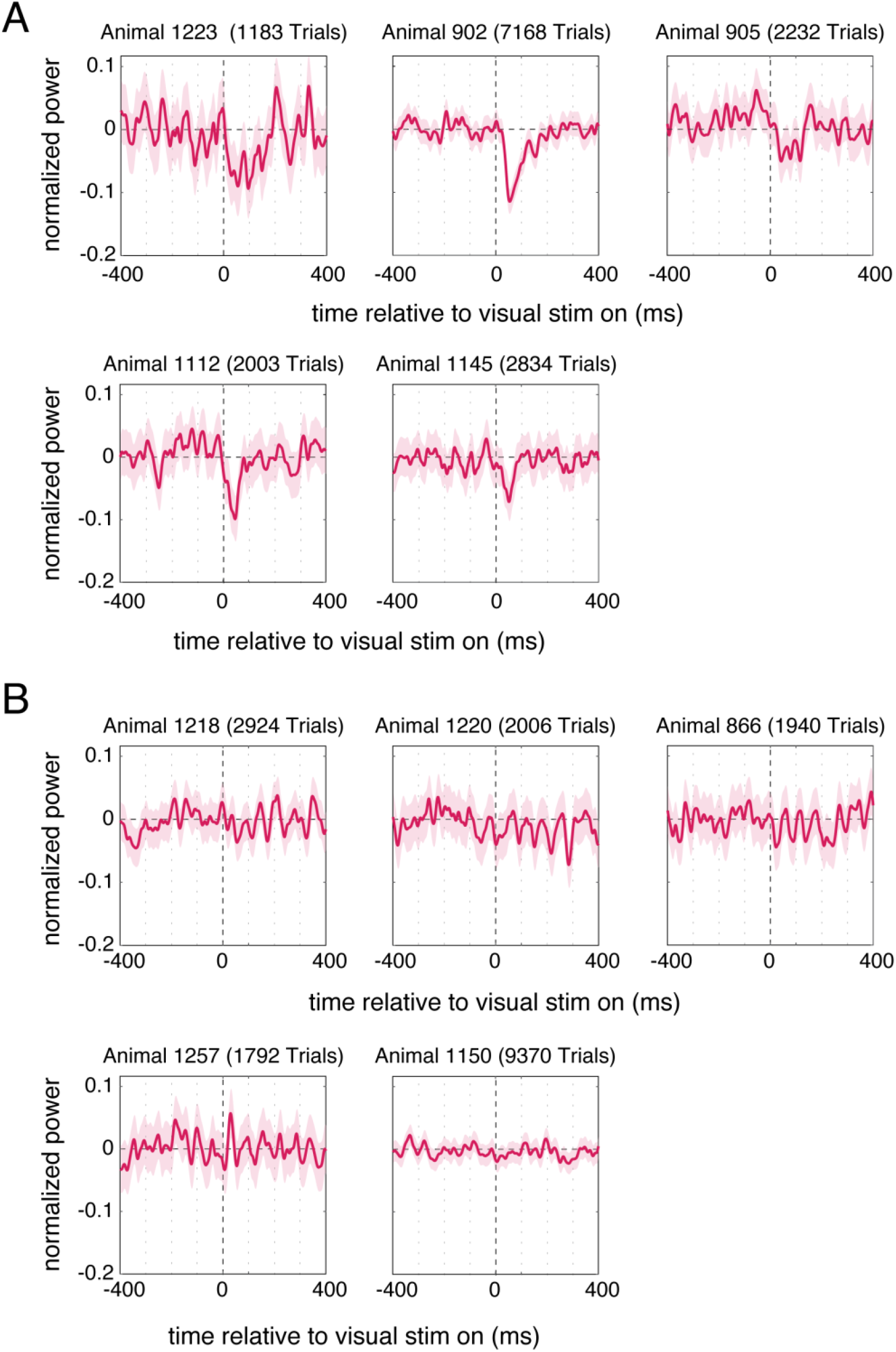
Neuronal-behavioral kernels for individual mice. Each plot represents the cumulative neuronal-behavioral kernel aligned to the onset of Gabor contrast steps for an individual mouse. **A)** Five mice showed significant modulations in the neuronal-behavioral kernel immediately after the onset of visual contrast steps. The shaded areas represent the 95% CIs (bootstrapped). **B)** Five mice did not individually have a clear modulation in the neuronal-behavioral kernel in the time surrounding the contrast steps. Variance in the magnitude of optogenetic effects likely arises from factors such as the strength of opsin expression over the course of data collection and number of neurons effectively illuminated by the fiber optic. Conventions are the same as in A.

**Figure S2.**
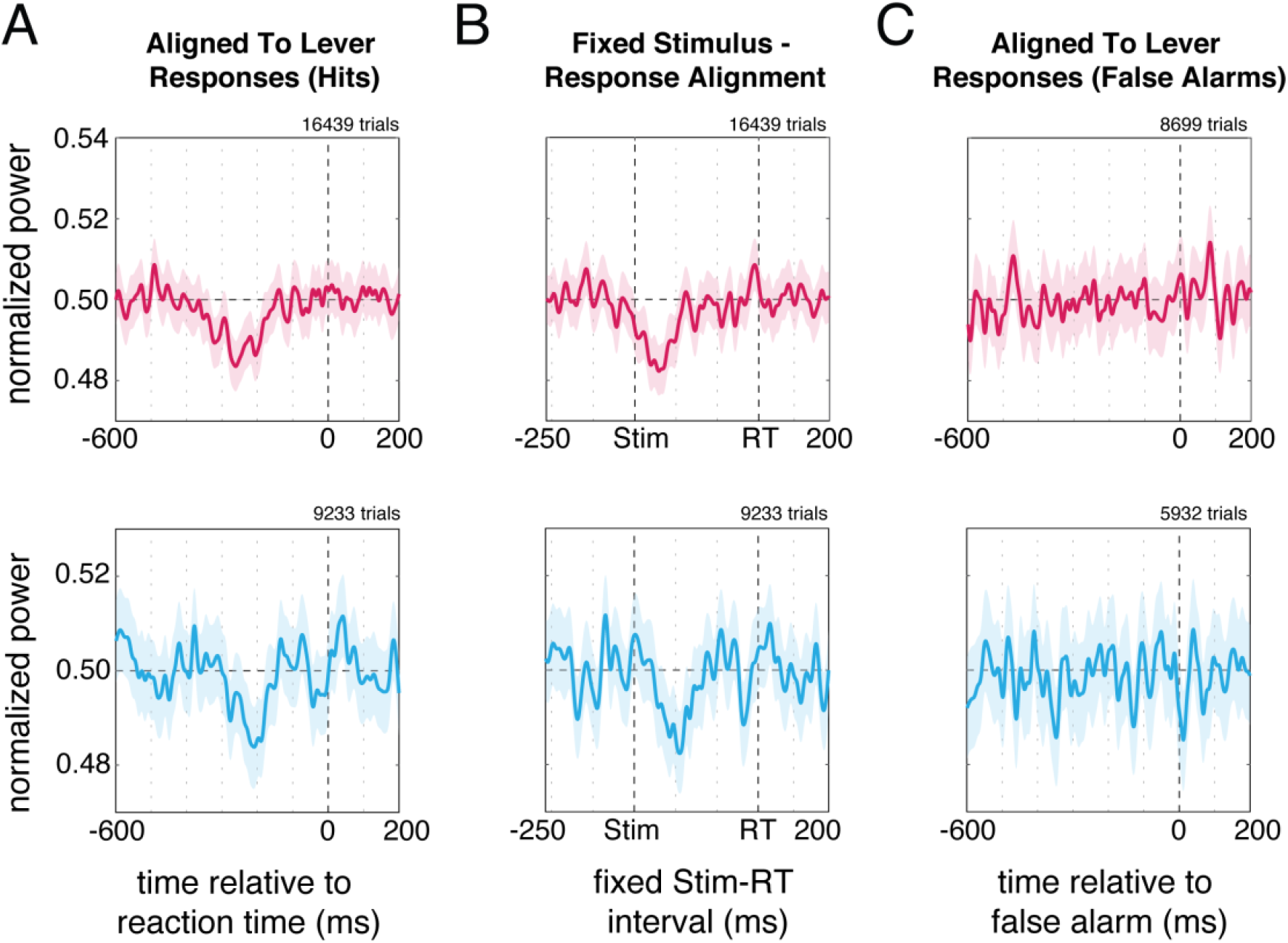
Neuronal-behavioral kernels aligned to behavioral responses. Behavioral kernels were aligned to behavioral responses for contrast steps (top row, red) and ramps (bottom row, blue). **A)** Optogenetic stimuli from hit trials aligned to lever releases. Significant modulations in the behavioral kernels occur hundreds of milliseconds before lever responses, at a time corresponding approximately to when the visual stimulus appeared. The greater breadth of these kernels compared to kernels that were aligned to the onset of visual stimuli (shown in Figure 2B, C) arises because the stimulus-response interval varies from trial to trial and the V1 contributions are better aligned to visual stimulus onset than to response times. **B)** Neuronal-behavioral kernels constructed using optogenetic stimuli that were stretched or compressed in time to bring the stimulus-to-response interval to 300 ms and thereby align all trials on both the stimulus onsets and the response times. A significant modulation is seen shortly after stimulus onset. The fact that these kernels are broader than the stimulus-aligned kernels (Figure 2) and narrower than the response time kernels (Figure S2A) supports the idea that V1 contributions are better aligned to visual stimulus onset than later stages of the perceptual/behavioral cycle. **C)** Behavioral kernels constructed by aligning optogenetic stimuli to lever releases on false alarm trials. The absence of significant modulations in the behavioral kernel argues that the false alarms were not generated by optogenetic stimulation producing a fictive perception of a stimulus.

**Figure S3.**
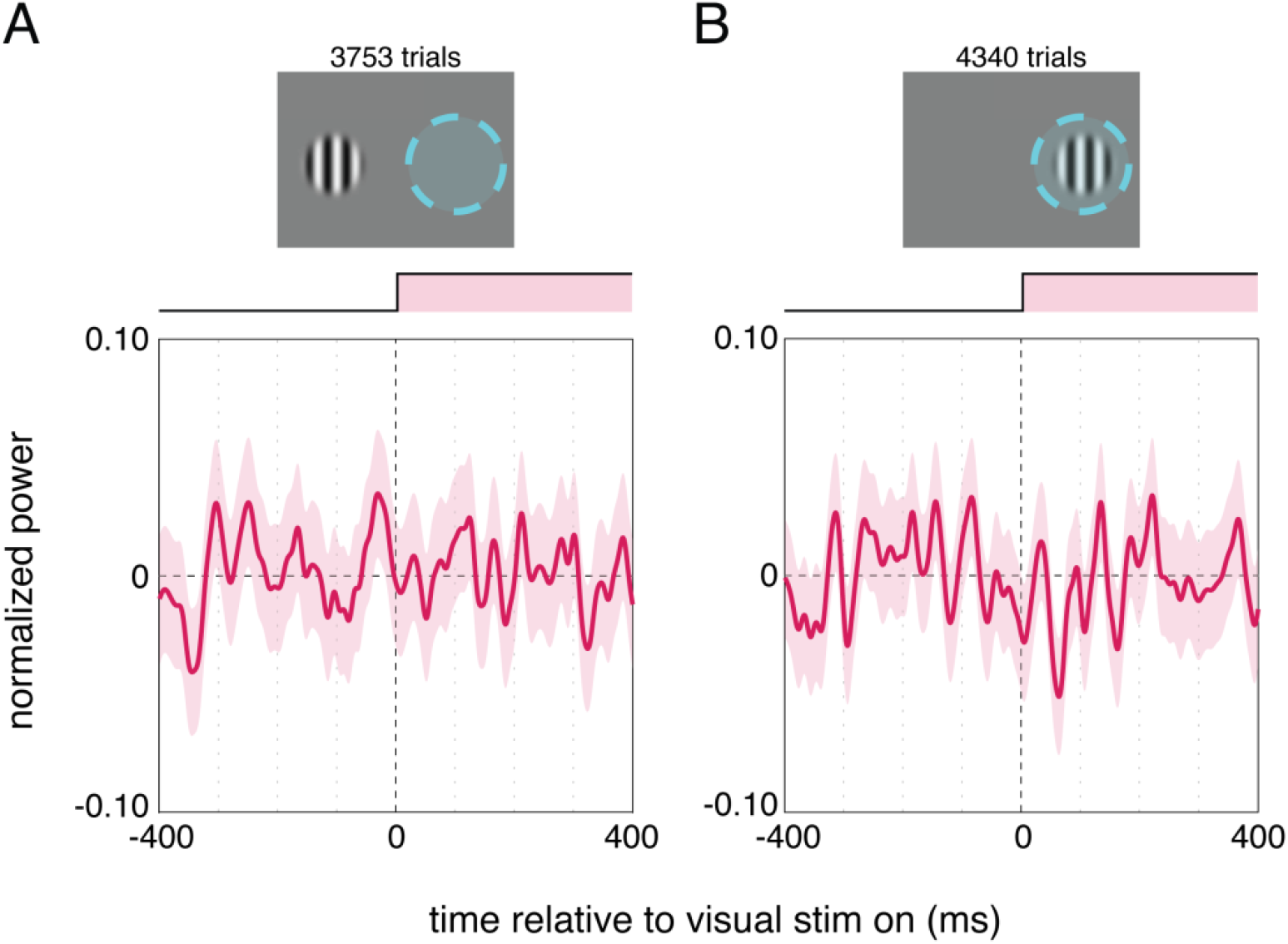
Behavioral effects of optogenetic stimulation depend on retinotopic alignment between visual and optogenetic stimuli. In control experiments, data were collected from 3 mice in which the retinotopic location of the visual contrast steps were either misaligned (A) or aligned (B) with the retinotopic location of optogenetic stimulation (see Methods). The red traces depict the cumulative neuronal-behavioral kernels and 95% CI. **A)** When visual and optogenetic stimuli are misaligned, there are no identifiable periods with significant modulations in the neuronal-behavioral kernel. **B)** When the site of optogenetic inhibition perturbs the patch of V1 that encodes visual stimulus, a significant modulation in the neuronal-behavioral kernel is observed during the initial 100 ms following visual stimulus onset. These data show that the effect of optogenetic stimulation is specific to the patch of V1 that represents the visual stimulus and that the effects of optogenetic stimulation on behavior did not arise from cortical heating due to light delivery, direct retinal stimulation, or other nonspecific effects.

